# Opportunities and challenges for applying Key Biodiversity Areas Criterion E at large spatial scales

**DOI:** 10.1101/2025.10.16.682361

**Authors:** Giordano Mancini, Marta Cimatti, Marianne Tzivanopoulos, Wilfried Thuiller, Moreno Di Marco

**Author notes:** Corresponding author: Giordano Mancini.

## Abstract

Key Biodiversity Areas (KBAs) are a cornerstone of global biodiversity conservation, influencing international strategic plans and helping protect thousands of species. KBAs are identified through quantitative criteria, among which the most recent is Criterion E. KBA Criterion E uses Spatial Conservation Prioritization techniques to identify highly irreplaceable sites, representing a promising tool for effective expansion of the KBA network. However, it has rarely been tested or applied at large scales. Here, we carried out a continental application of KBA Criterion E in Europe, using Species Distribution Models (SDMs) for 5,529 species of insects and 972 tetrapods. We stress-tested the application of Criterion E by changing the following settings: irreplaceability threshold, metrics of irreplaceability, representation targets, spatial resolution, and cost of planning units. Under the standard Criterion E settings, we identified 23 potential KBAs for insects, mostly along northern European coasts, and 88 for tetrapods, mostly concentrated in Mediterranean islands and southern Europe. These sites slightly overlapped with existing KBAs, showing that Criterion E can capture biodiversity patterns overlooked by other criteria. Our results also showed that the identification of highly irreplaceable areas is very sensitive to analytical choices. The strict irreplaceability threshold currently required, associated with the definition of representation targets, limited the selection of important sites almost exclusively to those containing very narrow-range species, and when such species were absent, important sites were preferentially selected on coasts, where the cost of planning units (represented by land extent) was minimized. Our analysis showed both opportunities and challenges of Criterion E and its applications with SDMs. We propose potential adjustments to the definition and guidelines of Criterion E, to improve its applicability at large spatial scales and on different taxa. Improvements of KBA Criterion E will ensure that KBAs continue to substantially contribute to the global conservation of biodiversity.

## Introduction

Key Biodiversity Areas (KBAs) are “sites contributing significantly to the global persistence of biodiversity” (IUCN, 2016). More than 16,500 KBAs have already been mapped worldwide, contributing to the representation of more than 18,000 species of conservation concern (BirdLife International, 2025b). KBAs are a useful tool to guide the expansion of protected areas (Kullberg et al., 2019) and can influence international strategic plans, for example, helping reach the goals set in the Kunming-Montreal Global Biodiversity Framework (Watson et al., 2023). Therefore, it is essential to improve the consistency and reliability of the identification of KBAs. Under the latest Global Standard (IUCN, 2016), KBAs are identified through five standardized criteria built upon quantitative thresholds, that are conceptualized to be applicable to very different taxa, ensuring consistency for the identification of important biodiversity areas. The criteria measure different biodiversity elements such as the presence of threatened or restricted-range species or ecosystems (Criteria A and B), high levels of ecological integrity (Criterion C), presence of important biological processes (Criterion D), or high levels of irreplaceability measured through quantitative analysis (Criterion E). Yet, while some of the criteria (such as that on threatened species) derive from decades-long use in previous KBA-like standards, Criterion E still requires substantial testing especially for large-scale applications (Smith et al., 2019).

Despite their crucial role in biodiversity conservation, some limitations have been historically highlighted in the KBA framework. In terms of methodology, the number of potential KBAs has been shown to increase with the number of species used to identify them, which could theoretically lead to an excessively high number of potential KBAs being identified (Farooq et al., 2023). Inherent inconsistencies also exist in the relationship between old and new KBA standards, for example 12,000 Important Bird Areas (IBAs; BirdLife International, 2025a) were converted into KBAs, although they have been identified using partially different criteria than those in the new Global KBA Standard. This led to a taxonomical bias in representation, as 69% of the current KBAs are identified using birds and plants (BirdLife International, 2025b), suggesting that KBAs are not yet representative of all biodiversity. In that context, building on the combination of extensive tools from Spatial Conservation Prioritization was considered as a potential solution to help overcome such limitations (Di Marco et al., 2016; Smith et al., 2019), allowing extensive and systematic evaluation of a large number of species over large spatial extents.

Spatial Conservation Prioritization is an integral part of Systematic Conservation Planning (Margules & Pressey, 2000). It builds on the use of optimization algorithms to prioritize the identification of a system of areas that efficiently achieves predefined representation targets, in order to better allocate resources for conservation (Moilanen et al., 2009). Spatial Conservation Prioritization is grounded in a number of key principles that need to be satisfied for effectively solving prioritization problems (Kukkala & Moilanen, 2013), such as complementarity, the degree of which a site adds previously unrepresented features to an existing network of areas (Margules & Pressey, 2000), efficiency, the ratio of benefits to costs in selecting certain areas (Kukkala & Moilanen, 2013), and irreplaceability. Irreplaceability was firstly defined by Pressey et al. (1994) as: “(i) the potential contribution of any site to a reservation goal; and (ii) the extent to which the options for a representative reserve system are lost if that site is lost.” Irreplaceability is a key concept for the KBA framework in the new Global Standard, as it provides the foundation for KBA Criterion E, which highlights sites as irreplaceable due to their distinct combination of biodiversity features using Spatial Conservation Prioritization approaches.

A site is eligible as a KBA under Criterion E if it “has a level of irreplaceability of ≥0.90 (on a 0–1 scale), measured by quantitative spatial analysis, and is characterised by the regular presence of species with ≥10 reproductive units known to occur (or ≥5 units for EN or CR species)” (IUCN, 2016). KBA Criterion E takes advantage of spatial prioritization tools to estimate the irreplaceability of sites (or planning units) and apply quantitative thresholds to identify potential KBAs. This requires specifying several settings prior to the analysis, such as the desired representation targets for each biodiversity feature and the spatial scale of analysis. The distribution of biodiversity features (i.e., species) used to apply KBA Criterion E can be represented in a number of ways including range maps, Area of Occupancy (AOO), or habitat suitability maps derived from Species Distribution Models (SDMs). SDMs are statistical models that relate species occurrences with a set of environmental variables to derive species niche and eventually project it geographically to predict species potential suitable habitat (Guisan et al., 2017; Thuiller, 2024). SDMs are used for a variety of purposes in conservation biology, from predicting species extinction risk (Mancini et al., 2024; Thuiller et al., 2011) to informing prioritization analyses (Araújo et al., 2011; Chauvier-Mendes et al., 2024; Guisan et al., 2013; Tulloch et al., 2016). SDMs are also a promising tool for the application of KBA Criterion E, delivering a better representation of species distribution compared to geographic ranges and being more readily available over large extents and a AOO for large numbers of species. Yet, the usability of SDMs in the context of KBA Criterion E remains to be comprehensively tested.

Although the Global KBA standard has been developed through a series of workshops, with the contribution of many experts from academia, conservation organizations and governments, synthesizing their expertise in the definition of KBA criteria (IUCN, 2016), there are still biases in the application of such criteria. For example, KBA Criterion E is one of the least used criteria, with just 207 KBAs (0.01% of all KBAs) identified, all which are in South Africa (BirdLife International, 2025b). Moreover, despite prioritization analyses linking irreplaceability to threshold-based criteria having been conducted for IBAs (Di Marco et al., 2016), a multi-taxa, continental, application of KBA Criterion E has never been done. Importantly, the potential effects of the various Criterion E settings have never been tested. Here, we applied KBA Criterion E using 5,529 insects and 972 tetrapods in Europe. We represented each biodiversity feature using SDMs and estimated irreplaceability using multiple metrics and under different settings. Our objectives were to: i) test the use of SDMs for identifying KBAs under Criterion E at a large scale and ii) to investigate how changing some of the settings required for the application of KBA Criterion E influenced the identification of potential KBAs. Based on our results, we provide some recommendations for future improvement of KBA standards and guidelines.

## Methods

We used SDM-derived habitat suitability maps of 5,529 European insects and 972 European tetrapods to estimate the irreplaceability of areas in Europe and apply KBA Criterion E. We estimated irreplaceability separately for insects and tetrapods. We then stress-tested KBA Criterion E changing several settings required for its application: the threshold to identify highly irreplaceable areas, the metric to represent irreplaceability, the representation targets, the spatial resolution of analyses, and the use of an areal “cost”.

### Species distribution data

We used habitat suitability maps of 5,529 European insects: 3,259 Heterocera (Lepidoptera), 475 Anthophila (Hymenoptera), 442 Carabidae (Coleoptera), 410 Syrphidae (Diptera), 344 Rhopalocera (Lepidoptera), 276 Orthoptera, 200 Ichneumonoidea (Hymenoptera), 115 Odonata, 108 Formicidae (Hymenoptera). Overall our insect data included 23% described European species for the considered groups (Tab. S1). We also used habitat suitability maps for 1,207 European tetrapods: 526 birds, 296 mammals, 268 reptiles and 117 amphibians. These represent ca. 73% of all European tetrapods (IUCN, 2023).

Habitat suitability maps for tetrapods and insects were produced through the European NaturaConnect project. Tetrapods maps are available from the EBV Data portal (Si-Moussi & Thuiller, 2024) and insect maps from (Tzivanopoulos et al., in review). Models were calibrated using data obtained mainly from GBIF (GBIF.org, 2023), as well as IUCN and BirdLife (for tetrapods), and observation.org and the Safeguard pollinator database (for insects). Observations were cleaned using the ‘CoordinateCleaner’ R package (Zizka et al., 2019). Although an IUCN range was available for many tetrapods to generate true absences, insect data was presence-only, therefore background points were produced using a target-group approach accounting for sampling effort. Models were calibrated using climatic (CHELSA; Brun et al., 2022; Karger et al., 2017), soil (SoilGrids; Poggio et al., 2021) and land-use predictors (Sandström et al., 2023) for all groups, with some additional hydrology and geology variables (GliM, EU-Hydro) used in tetrapod models. An ensemble of machine learning methods was trained including Random Forests and XGBOOST (both groups), a Multi-Layer Perceptron neural network (tetrapods), Generalized Linear and Additive models (insects). Model calibration was done with spatial-block cross-validation over the study area and parts of northern Africa and western Asia to avoid niche truncation. Species habitat suitability was projected using Community Averaging algorithms integrating model uncertainty and applied the Estar framework to constrain model predictions based on observation density, biogeography and IUCN expert range when available.

Habitat suitability maps were produced at a ∼1km resolution in Lambert Azimuthal Equal Area (LAEA) Europe projection. We excluded all species with <10% of the suitable habitat in the study area (KBA Standards and Appeals Committee of IUCN SSC/WCPA, 2022), which encompasses continental Europe, excluding Belarus, Russia and Ukraine, resulting in a final sample of 5,529 insects and 972 tetrapods.

### Data preparation

We divided Europe into square cells of 10×10 km resolution, which complies with the resolution required by the Global KBA standard to represent planning units (PUs) in the spatial prioritization analysis for the application of KBA Criterion E (IUCN, 2016). We assigned a “cost” to each PU based on the amount of land in that PU (as per Global KBA standard; IUCN, 2016). For each PU, we calculated the amount of suitable habitat for each species (i.e., the prioritization features) whose habitat intersected that PU. We then defined the representation target for each species following the Global KBA standard (Tab. 1). The representation targets are defined based on the proportion of species’ habitat within the study area. For example, if a species had part of suitable habitat outside the study area, the representation targets are rescaled based on the proportion of suitable habitat within the study area to reflect the fraction of the species’ global population size in the study area (Tab.1; KBA Standards and Appeals Committee of IUCN SSC/WCPA, 2022).

**Tab. 1.** Representation targets for prioritization analysis under KBA Criterion E.

| Species' whole habitat within study area | Species' habitat also outside study area |
| --- | --- |
| if distribution $\geq 1,000 \text{ km}^2$:<br>Target = $\max(10\% \text{ distribution size, } 1,000 \text{ km}^2)$<br>if distribution $< 1,000 \text{ km}^2$:<br>Target = whole distribution size | if total distribution $\geq 1,000 \text{ km}^2$:<br>Target = $\max(10\% \text{ distribution within the study area, } 1000 \text{ km}^2 \times \frac{\text{distribution within study area}}{\text{total distribution}})$<br>if total distribution $< 1,000 \text{ km}^2$:<br>Target = whole distribution within the study area |

### Irreplaceability analysis and application of KBA Criterion E

We performed separate prioritization analyses for insects and tetrapods and derived irreplaceability values of each individual grid cell in our study area. We performed the analyses using Marxan (Ball et al., 2009), one of the most used tools for spatial prioritization analysis, which is also suggested by the KBA guidelines for applying Criterion E (KBA Standards and Appeals Committee of IUCN SSC/WCPA, 2022). We used Marxan to calculate the selection frequency of each PU, which is a proxy of irreplaceability representing the frequency in which a planning unit is selected as part of a set of spatial solutions (i.e. following the definition of Pressey et al. (1994)). Marxan uses heuristic solvers such as the simulated annealing algorithm (Serra et al., 2020), which provides a number of near-optimal solutions estimated stochastically, where the selection frequency represents the proportion of times a PU is selected across all representative solutions (Serra et al., 2020).

Marxan needs to be calibrated to ensure the produced solutions are close to the lowest cost (Ardron et al., 2010). Due to the stochastic nature of the simulated annealing algorithm, the number of iterations will determine how many combinations could be explored to find the optimal set of solutions. The Species Penalty Factor (spf; or Feature Penalty Factor) should also be calibrated as it controls the penalty added to the objective function if the target for a conservation feature is not met (Ardron et al., 2010). The higher the spf the more emphasis will be placed ensuring that feature’s target is met. We set 100 runs for Marxan and iteratively tested all possible combinations setting the number of iterations to 100,000, 1,000,000, 10,000,000 and 100,000,000 and the sfp to 100, 200, 300, 500, 600, 1000, 5000.

We performed the calibration for both insects and tetrapods at 10 km and 30 km resolution (see below) and selected the run that minimized the cost needed for the best set of solutions. The final Marxan settings for insects at 10 km resolution were 100,000,000 iterations and a spf of 200; while the settings at a 30 km resolution were 100,000,000 iterations and a spf of 5000 (Appendix S1). The final Marxan settings for tetrapods at 10 km resolution were 100,000,000 iterations and a spf of 300; while the settings at a 30 km resolution were with 100,000,000 iterations and a spf of 500 (Appendix S1).

We applied KBA Criterion E separately for insects and tetrapods to find potential European KBAs for each group. A site qualified as KBA under KBA Criterion E if it “has a level of irreplaceability of ≥0.90 (on a 0–1 scale), measured by quantitative spatial analysis, and is characterised by the regular presence of species with ≥10 reproductive units known to occur (or ≥ 5 units for EN or CR species)” (IUCN, 2016). We scaled the 0-100 values of selection frequency to 0-1 matching the scale required by KBA Criterion E. Then, we selected all PUs for which selection frequency ≥0.9. We then used occurrences for each species, from the Global Biodiversity Information Facility (GBIF), to account for potential presence of reproductive units. We considered confirmed presence of species when at least one occurrence point falls within the highly irreplaceable PUs.

### Stress testing the application of KBA Criterion E

To test the functionality and reliability of KBA Criterion E, we performed additional analyses calculating the irreplaceability changing all settings required by the Global KBA standard, one at a time, while keeping all the rest as standard settings. In particular we tested: the threshold to identify highly irreplaceable areas, the metric to represent irreplaceability, the representation targets, the spatial resolution of analyses, and the use of an areal “cost” (Tab. 2).

**Tab. 2.** Summary of the settings tested for KBA Criterion E application.

| Parameters | Settings |
| --- | --- |
| Irreplaceability threshold | 0.5, 0.7, 0.9 |
| Irreplaceability metrics | Selection frequency, Ferrier's importance, replacement cost |
| Resolution of planning units | 10 km, 30 km |
| Cost of planning units | Areal cost, unitary cost |
| Representation targets | Real target, target x2, target x3 |

#### Threshold level of irreplaceability

We tested how the number of PUs selected as highly irreplaceable changed for different thresholds of irreplaceability. We compared the number of PU selected using thresholds of selection frequency ≥0.5 and ≥0.7 with those selected using the standard settings (i.e., selection frequency ≥0.9). This test was done to check if the current threshold of irreplaceability was too restrictive in terms of PUs selected as potential KBAs.

#### Irreplaceability metrics and prioritization software

In addition to the selection frequency, we calculated two other metrics that approximate irreplaceability: “Ferrier’s importance”, a statistical estimator of irreplaceability (Ferrier et al., 2000), and “Replacement Cost”, which refers to “the increase in cost needed to achieve the targets after an inclusion/exclusion of a site (or set of sites)” (Cabeza & Moilanen, 2006). Ferrier’s importance is calculated for each site and for each single feature (Ferrier et al., 2000). We considered the “summed irreplaceability” which is calculated summing irreplaceability values associated to all sites with the contribution of each feature, as proposed by Ferrier et al. (2000), which potentially ranges from 0 to the number of features considered. We also used the maximum value obtained for each PU (regardless of the feature) as our other metric representing Ferrier’s importance (named here Relative max Ferrier’s importance) which ranges between 0 and 1. The replacement cost already provides a single value for each PU. A replacement cost value of 0 means that alternative solutions with the same properties exist, while values larger than 0 mean that any alternative solution including/excluding the focal site (i.e., PU) will have a higher cost than the optimal one (Cabeza & Moilanen, 2006). To calculate Ferrier’s importance and replacement cost we used the ‘prioritizr’ package, that takes advantage of different solvers to perform mixed integer linear programming providing a mathematically optimal solution (Hanson et al., 2024). We used the Gurobi solver, which was found to be the best in terms of computational speed for large scale problems (Hanson et al., 2024). We set the objective to minimize the cost of the best solution (i.e., minimum set problem), as done in Marxan. We used the same settings and input data used to calculate the selection frequency (i.e., the standard settings required for the application of KBA Criterion E).

#### Spatial resolution of planning units

The Global KBA standard requires prioritization analysis to run on PUs with a spatial resolution of 100-1,000 km^2^ (IUCN, 2016). Thus we repeated the selection frequency analysis for insects and tetrapods using the standard settings described above, but this time using PUs of 30×30 km (900 km^2^) resolution. This test was done to identify potential changes in the selection or composition of systems of highly irreplaceable PUs due to changing resolution.

#### Cost of planning units

Global standards for the application of KBA Criterion E require setting the cost of each PU as the proportional extent of area in that PU (KBA Standards and Appeals Committee of IUCN SSC/WCPA, 2022). As our analysis focuses on terrestrial species, the proportion of area in each PU corresponds to the proportion on land, meaning that PUs on the coasts should have lower cost than PUs inland. This could affect the output of prioritization softwares designed to resolve a minimum set problem, where the goal is to achieve representation targets at the smallest possible cost (McDonnell et al., 2002), and could lead to preferences for coastal PUs (“cheaper” PUs). Therefore, we tested the potential impact of the PUs’ cost by calculating the selection frequency setting a unitary cost for all PUs (i.e., cost=1 for all PUs) and comparing the results with those obtained with standard settings (i.e., area extent cost).

#### Representation targets

Finally, we tested how the selection of highly irreplaceable PUs could change based on different representation targets. We compared PUs selected using standard targets with those obtained doubling and tripling the standard targets (Tab. 2). For example, when doubling-up the target if the whole habitat was within the study area and ≥2,000 km^2^ we used *max(20% distribution, 2,000 km*^*2*^) or the whole distribution if it was <2,000 km^2^.

All analyses were performed using Marxan (Ball et al., 2009), Gurobi (Gurobi Optimization, 2024) and R 4.4.3 (R Core Team, 2025) in RStudio 2023.6.0.421 (Posit Team, 2023), with the following packages: ‘data.table’ (Barrett et al., 2024), ‘patchwork’ (Pedersen, 2023), ‘prioritizr’ (Hanson et al., 2024), ‘rgbif’ (Chamberlain & Boettiger, 2017), ‘terra’ (Hijmans, 2024), ‘tidyverse’ (Wickham et al., 2019).

## Results

### Irreplaceability analysis and application of KBA Criterion E

Using Marxan and the standard KBA settings at a 10 km resolution, we found 23 highly irreplaceable PUs (selection frequency ≥0.9) for insects and 88 highly irreplaceable PUs for tetrapods (Fig. 1). Interestingly, these PUs were located in opposite regions of the study area. Except for one irreplaceable PU in South Spain, all highly irreplaceable PUs for insects were in Northern Europe, on coasts of the Netherlands, Germany, on the Bornholm island (Denmark) and one on the southern Swedish coast (Fig. 1), while all highly irreplaceable PUs for tetrapods were in Southern Europe on Greek islands, Malta, Cyprus and in central Spain (Fig. 1). Importantly, the selection of 80 out of the 88 (90%) highly irreplaceable PUs for tetrapods was driven by the presence of four very restricted range species, for which the representation target was set equal to their entire distribution: *Mediodactylus oertzeni, Plecotus gaisleri, Podarcis filfolensis* and *Podarcis levendis*. None of the insect species had a distribution smaller than 1,000 km^2^ based on SDMs predictions. For both groups, the vast majority of PUs had low selection frequency value between 0 and 0.25 (Fig. 1). However, we found a clear coastal pattern of irreplaceability, where PUs on the coasts had higher selection frequency values than PUs inland in both analyses due to their lower cost in terms of land-area surface (Fig. 1).

**Fig. 1.**
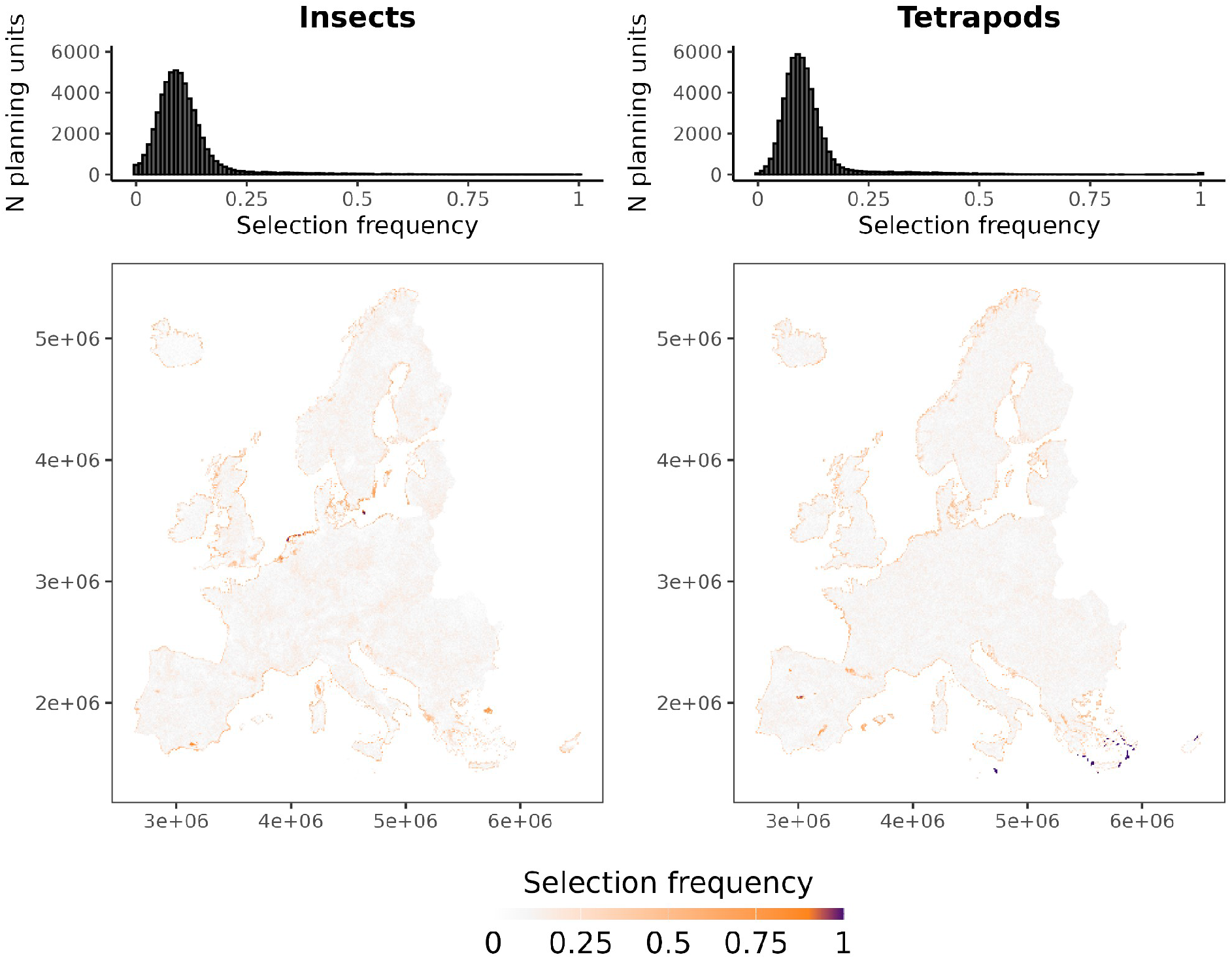
Selection frequency values at 10 km resolution. Maps represent the geographical pattern of selection frequency values for insects and tetrapods. Histograms represent the distribution of selection frequency values for insects and tetrapods.

Although many insect and tetrapod species had >100 known occurrences in GBIF (Fig. S1a,c), a large proportion of them did not have any occurrence falling in the highly irreplaceable PUs (Fig. S1b,d). However, we found 1,631 insect species and 98 tetrapod species with at least 1 occurrences (representing confirmed presence here) within the highly irreplaceable PUs found for each group.

### Stress testing the application of KBA Criterion E

When changing the irreplaceability threshold to identify irreplaceable PUs we found similar results across insects and tetrapods, with 596 and 586 PUs selected under a selection frequency threshold of 0.5, and 101 and 130 PUs under a threshold of 0.7 respectively (Fig. 2). When exceeding the selection frequency threshold of 0.7, we found a sharp decline in PUs identified for insects, while the number of PUs identified for tetrapods remained almost constant.

**Fig. 2.**
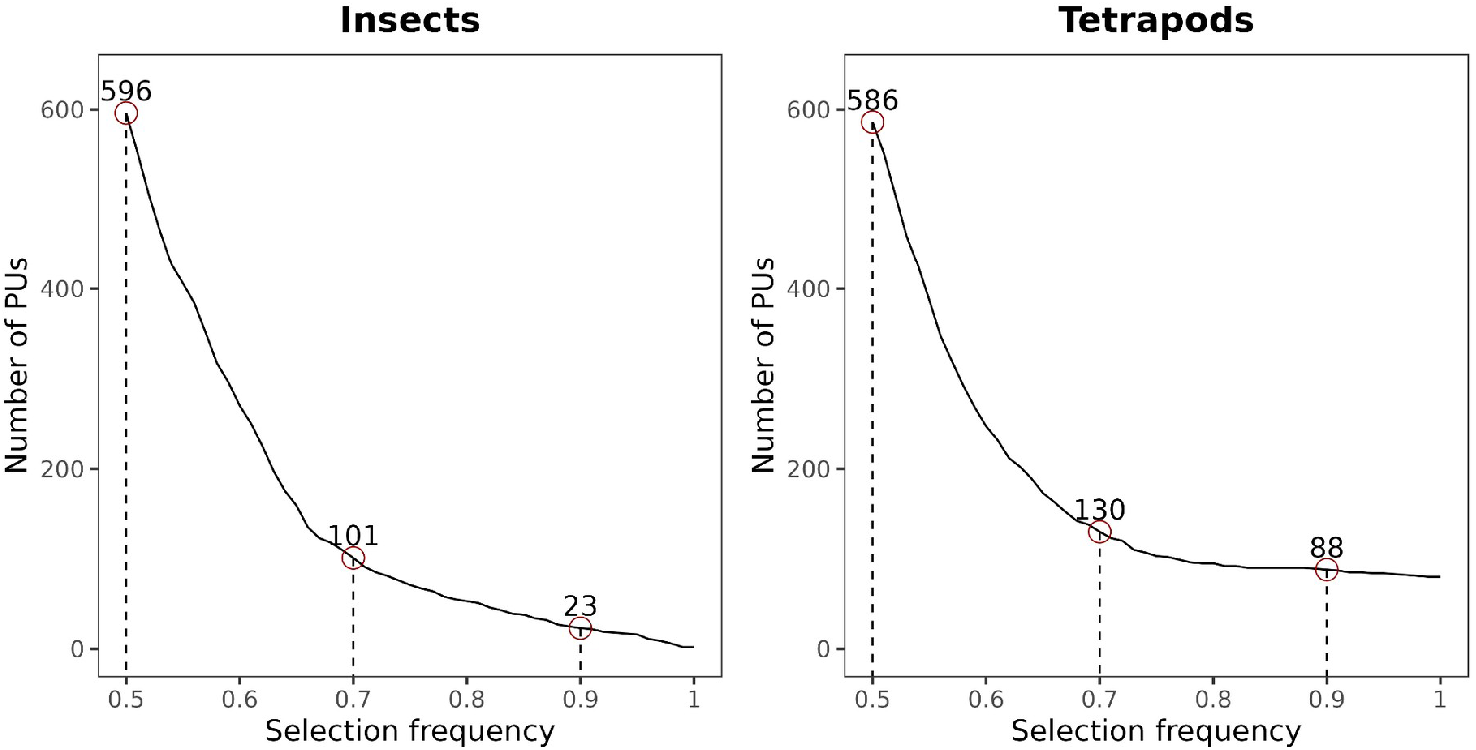
Number of planning units (PUs) selected using different thresholds of selection frequency for insects and tetrapods. Red circles and numbers represent the number of PUs identified at selection frequency thresholds of 0.5, 0.7 and 0.9.

We compared values of Ferrier’s importance and replacement cost with the selection frequency calculated for insects and tetrapods. For insects, we found 13 highly irreplaceable areas identified by the replacement cost (Fig. 3) and none by the Ferrier’s importance (both metrics used; Fig. 3, S2). None of these PUs were in common with those identified by the selection frequency (k=0.08 between selection frequency and replacement cost). PUs identified by the replacement cost were all in central Europe (between France and Germany) and none of them close to a coast (Fig. 3). For tetrapods, we found 84 highly irreplaceable PUs identified by the replacement cost and 80 PUs identified as highly irreplaceable by the Ferrier’s importance (both metrics; Fig. 3, S2). We found 80 common PUs between all the metrics considered (k=0.95 between selection frequency and relative max Ferrier’s importance, k=0.93 between selection frequency and replacement cost and k=0.97 between relative max Ferrier’s importance and replacement cost), they are all the PUs where the four very narrow range species were present.

**Fig. 3.**
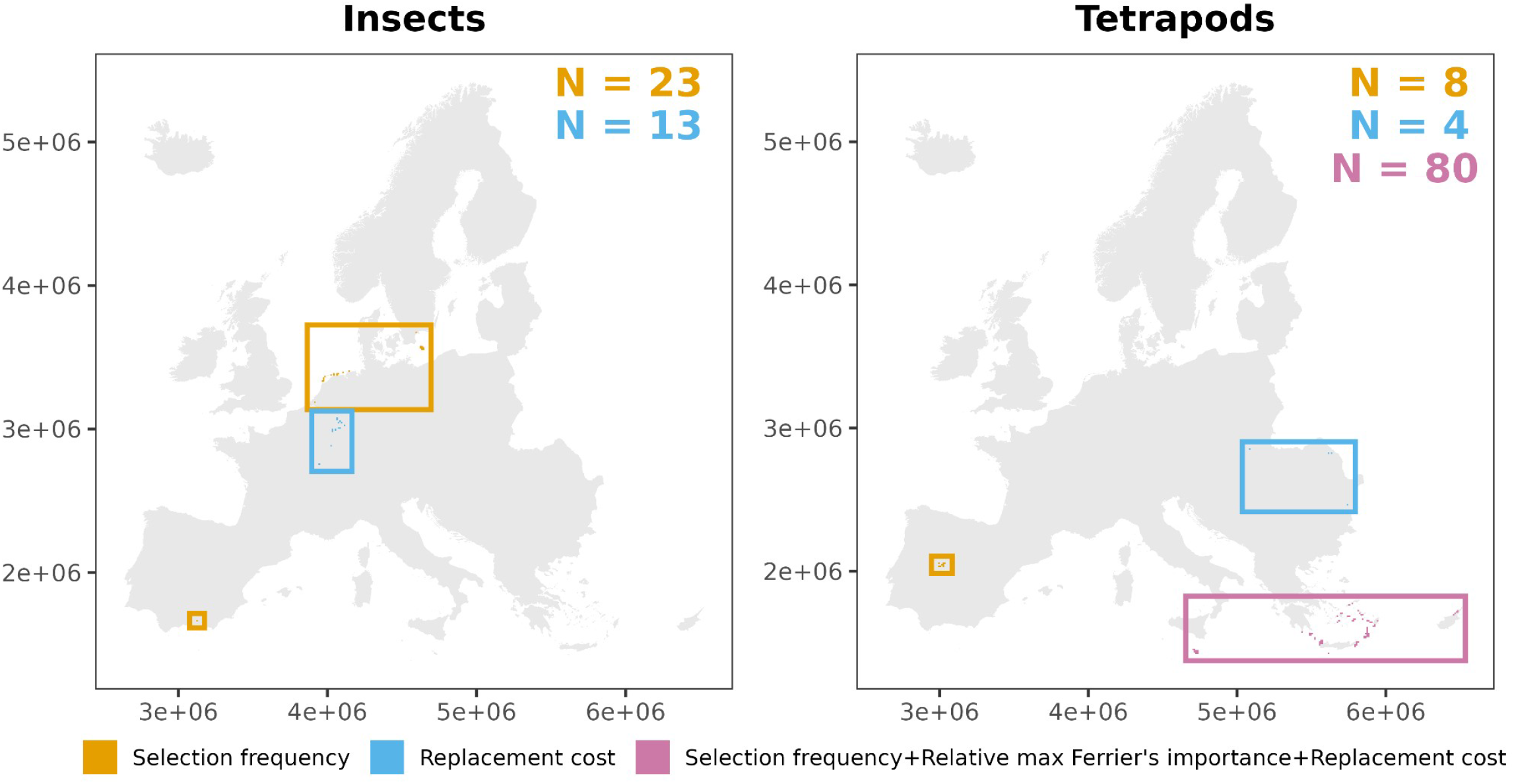
Highly irreplaceable planning units (PUs) at 10 km resolution identified by the selection frequency (orange), replacement cost (blue) or by all metrics (selection frequency, replacement cost and relative max Ferrier’s importance; red) for insects and tetrapods. Rectangles encompass highly irreplaceable PUs identified by the metric corresponding to the color. N: number of PUs.

We found a common pattern between the analyses performed at 10 km resolution and those at 30 km resolution for both insects and tetrapods (Fig. S3). We found 12 highly irreplaceable PUs for insects and 46 for tetrapods. For both groups the spatial distribution of the selection frequency values were very similar to the one obtained at 10 km resolution. In particular, we found that the highly irreplaceable PUs at 30 km resolution encompassed 19 of the 23 PUs found for insects at 10 km resolution and 87 of the 88 found for tetrapods.

When changing the cost of PUs to a unitary value we found much lower irreplaceability in coastal areas compared to the original analysis (Fig. S4a,b). This confirms that intermediate irreplaceability was mainly driven by cost (i.e. land-area surface) rather than species content under our analytical settings. We also found some differences for PUs with very high irreplaceability between insects and tetrapods. For insects, we found 26 highly irreplaceable PUs with analysis at unitary cost distributed across the Netherlands, Greece and Spain, with 10 of them in common with the analysis at standard cost (where each PU has a cost representing the extent of land in that PU; Fig. S4c). Overall, the analysis at standard cost selected PUs where 52% of the site was covered by land on average, in contrast to the average 91% of the PUs selected at unitary cost: the analysis at standard costs preferred PUs on the coastline. For tetrapods, the analysis at unitary cost identified 88 highly irreplaceable PUs, and 87 of them were in common with the analysis at standard cost (Fig. S4d): the two analyses identified almost the same set of highly irreplaceable PUs.

As expected, increasing the target led to a greater number of PUs identified as irreplaceable. When we doubled the target, we found more than twice as many PUs for insects as for tetrapods (865 and 326 highly irreplaceable PUs respectively; Fig. 4), and when tripling it we found 1,219 and 1,303 PUs for insects and tetrapods. The geographical distribution of the selected PUs based on different targets resembled the one found for the standard target, with PUs for insects mostly located on the North European coasts (e.g., coasts of The UK, France, the Netherlands, Germany, Denmark and Sweden; Fig. 4) and those for tetrapods overall equally distributed across South Europe (mostly Spain, Italy and Greece; Fig. 4).

**Fig. 4.**
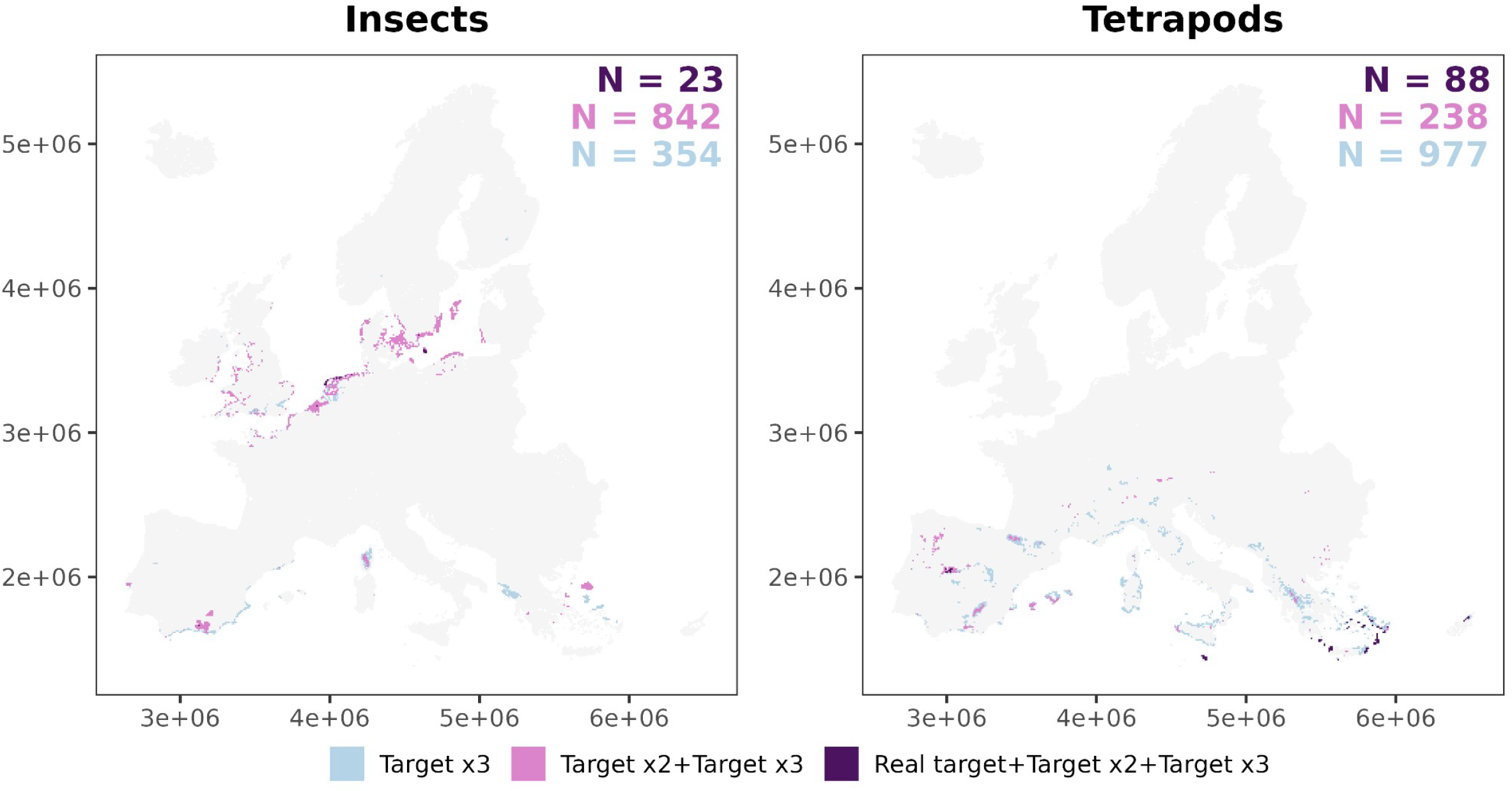
Highly irreplaceable planning units (PUs) at 10 km resolution identified by different targets for insects and tetrapods: tripled target only (lightblue), tripled and doubled target only (pink), or tripled, doubled and original target (purple). N: number of PUs.

## Discussion

### Potential Key Biodiversity Areas under Criterion E

We found 23 potential KBAs for insects and 88 for tetrapods in Europe under current KBA settings, using SDMs to represent biodiversity features at a European continental scale. The newly identified sites were located in different areas, largely non-overlapping: insect sites were concentrated in northern Europe, especially along the coasts of the Netherlands, Germany, Denmark, and southern Sweden, whereas tetrapod sites were located in southern Europe, particularly Mediterranean islands and parts of the Iberian Peninsula. This strong geographic contrast demonstrates that Criterion E can capture taxon-specific patterns of biodiversity that complement, but do not duplicate, those highlighted by existing KBA criteria. In fact, among existing KBAs, only 6 overlapped with insect KBAs and 54 with tetrapod KBAs, while simultaneous overlap across all three designations (existing KBA, insect KBA, tetrapod KBA) was absent. This mismatch is unsurprising: most existing European KBAs originate from Important Bird Areas and sites for threatened plants (BirdLife International, 2025b), which were identified through different criteria. This restricted correspondence suggests that KBA Criterion E provides additionality to the KBA network, especially for underrepresented taxa such as insects.

The differences in the identification of the important sites between insects and tetrapods were mostly caused by the presence of very narrow range species. Among tetrapods, four species (*Mediodactylus oertzeni, Plecotus gaisleri, Podarcis filfolensis, Podarcis levendis*) with very narrow distributions, drove the identification of almost all highly irreplaceable PUs. This reflects the intent of KBA Criterion E, which is designed to highlight globally unique biodiversity features (IUCN, 2016), and aligns with the findings of Di Marco et al. (2016), who showed that irreplaceability increases with the presence and abundance of restricted-range species. However, it also illustrates that irreplaceability outputs, in the context of KBA Criterion E, may be dominated by very few range-restricted species which have a disproportionate importance in the calculation of the selection frequency, as for such species the representation target has to be set as the species’ entire distribution (IUCN, 2016), potentially obscuring broader patterns. On the contrary, none of the insect species had a very narrow distribution, thus the selection frequency was calculated by exploring different combinations of PUs that reach all targets minimizing the cost (minimum set problem). This led to the identification of a set of few highly irreplaceable PUs, but many of these were located on coasts of the study area, where the cost, represented by the extent of land, was minimized.

The results we found for insects should be interpreted considering their taxonomical and geographical representation bias. Occurrence data, needed for SDMs, is lacking for insects, especially for endemic ones, leading to a bias towards common species (Troudet et al., 2017). For example, Orthoptera show very high species richness in Southern Europe, including several endemic species (Kenyeres et al., 2009), yet this region shows much less available data compared to Northern Europe, where some countries (e.g., the Netherlands and Sweden) have extremely good sampling schemes and many common species were better represented in the species pool used to produce habitat suitability maps for insects (Tzivanopoulos et al., in review). Such bias can affect irreplaceability analysis and should be accounted for when applying KBA Criterion E.

Despite these methodological sensitivities, the geographic distribution of highly irreplaceable sites corresponds well to known biodiversity patterns. For tetrapods, the concentration of KBAs in southern Europe aligns with established centers of endemism for reptiles, amphibians, and small mammals (Cox et al., 2006; Myers et al., 2000). For insects, the emergence of coastal North European sites as important areas is novel but ecologically plausible, reflecting unique assemblages of insects adapted to wetland, and island ecosystems. These regions are often poorly represented in existing conservation networks, suggesting that Criterion E can highlight overlooked biodiversity values.

Based on our results, we provide a number of recommendations either for the Global KBA standard and for the KBA guidelines to enhance the robustness and reliability of KBA Criterion E applications.

### Recommendations for the Global KBA Standard

Our stress-testing revealed several important sensitivities in applying Criterion E. We found that the majority of the PUs in both insect and tetrapod analyses had selection frequency between 0 and 0.25, sharply decreasing beyond those values. The high irreplaceability threshold of 0.9 currently required to identify potential KBAs, in association with the low representation targets of 10% of species distribution, represent an important constraint to the selection of highly irreplaceable areas. In our analysis, such a threshold was reached mostly by PUs where species with very narrow ranges occur, or by few PUs where many species overlap. We suggest that under current target formulation, irreplaceability threshold could be relaxed to some extent without risking excessively high numbers of KBAs to be identified. For example, using a threshold of 0.7 or 0.75 adds very little PUs compared to the 0.9 threshold currently required. This would allow capturing either species rich PUs, still with a high level of irreplaceability, along with very highly irreplaceable areas identified by the presence of species with very narrow range.

The overall very low levels of irreplaceability we found were probably also driven by the large distribution range of species in our dataset. Small targets generally reduce the level of irreplaceability, especially if the feature’s distributions are large, as only a few PUs are needed to reach the targets (i.e., the more overlap the distributions the fewer PUs are needed to reach the targets). Thus, a few species for which large targets are set (100% of distribution for restricted range species), will guide the final solution (Moilanen & Arponen, 2011). As currently formulated, the representation targets are too small to really guide the selection of irreplaceability in areas other than those with species with very narrow ranges.

We suggest considering expanding the representation targets. For example, doubling them up leads to many more, but still reasonable numbers of, irreplaceable PUs. These additional sites were also located in previously unrepresented, but important, regions, such as mediterranean mountains in Corsica for insects and Pyrenees and Alps for tetrapods, contributing to the representation of biomes of known importance for biodiversity (Körner, 2004).

Finally, we did not find major differences between analyses performed at 10 and 30 km resolution for insects and tetrapods: changing the resolution has little effect on the total selected area. We suggest testing for higher resolutions when regionally appropriate. However, this ultimately depends on the area under study (continental or national assessment) and on the available input data, as increasing the resolution could help handle commission errors (e.g., in the case of range maps) but risks selecting much more area than needed to identify potential KBAs (Di Marco et al., 2017), a criticism already pointed out for the KBA framework (Farooq et al., 2023).

### Recommendations for the Guidelines for using the Global KBA Standard

We used selection frequency, replacement cost and Ferrier’s importance to represent irreplaceability in our prioritization analysis. We found some consensus results between the metrics, along with some differences. All metrics identified almost the same set of highly irreplaceable areas for tetrapods. The PUs selected were those where very range restricted species occur and, as expected, all metrics assigned the highest irreplaceability values to such PUs. However, we found some differences in the selection of some PUs for tetrapods and all PUs for insects. Such differences were expected as among insects none had the largest target (100% of distribution), thus there is no constraint on the selection of certain PUs, and all metrics used are conceptually different. For example, the selection frequency measures how many times a PU is selected in a final set of solutions (Serra et al., 2020), while the replacement cost measures the increase in cost needed to achieve the targets if a PU is excluded (Cabeza & Moilanen, 2006). Importantly, the metrics were calculated by softwares that use different types of algorithms (heuristic vs exact algorithms). Therefore, the PUs selected by each metric have very different meanings, but all provide a measure of irreplaceability that complies with KBA Criterion E requirements (i.e., complementarity-based analysis). When applying prioritization analysis for KBA Criterion E, we suggest using at least two metrics of irreplaceability. The metrics will probably converge on selecting the very high irreplaceable PUs, but could also provide an additional set of irreplaceable PUs that should be considered and evaluated on a case by case basis.

We also recommend defining a proper scale for using the “summed irreplaceability” (Ferrier et al., 2000) to apply KBA Criterion E. As currently formulated, KBA Criterion E identifies sites with a level of irreplaceability of ≥0.9 on a 0-1 scale (IUCN, 2016). While the selection frequency and replacement cost can be provided in a scale that is easily convertible to a 0-1 scale (e.g., selection frequency usually ranges from 0 to 100), this is not the case for Ferrier’s importance. For Ferrier’s importance, each irreplaceability value associated to each site ranges between 0 and 1, depending on the features contributing to it, but the resultant “summed irreplaceability” can be >1 (Ferrier et al., 2000). One, in principle, could apply a simplistic scaling calculation, for example, dividing all irreplaceability values by the highest value, obtaining a scaled distribution between 0 and 1. However, this procedure could be misleading, because the presence of an exceptionally high value of irreplaceability (e.g. PUs hosting species with very restricted distributions) would cause all other values to be skewed, making them appear much lower than they actually are. In this study, we attempt to use a “relative max Ferrier’s importance” using the highest value of irreplaceability calculated for each site, which ranges from 0 to 1. This however conceptually deviates from the original formula proposed by Ferrier et al. (2000), as it relies on the highest contribution of a feature ignoring the contribution of the other ones, although many features contributing with modest irreplaceability values (e.g., 0.3) might still trigger high cumulative irreplaceability for a certain PU.

When testing for cost effect, we found a similar number of highly irreplaceable PUs for insects between the analysis with standard cost or unitary cost, but less than half of them (43%) were in common, while for tetrapods we found a large consensus of PUs (>90% in common). This consensus was almost entirely driven by the range-restricted species for which the representation target was the whole species’ distribution (see Tab. 1). This is in line with previous studies suggesting that the cost has very little effect on the selection of high priority PUs (Carwardine et al., 2010): sites where range restricted species occur are always selected to meet the representation targets, regardless of the cost assigned. However, the cost had an effect on the identification of highly irreplaceable PUs for insects, where the selection was not driven by the narrow range species. The KBA framework explicitly aims to identify sites that contribute significantly to the global persistence of biodiversity, not to prioritize sites for conservation action, thus no data on economic cost are considered (Smith et al., 2019). Instead a “cost” representing the extent of land in each PU is used in prioritization analysis to apply KBA Criterion E (KBA Standards and Appeals Committee of IUCN SSC/WCPA, 2022). Nevertheless, the use of such cost has an effect on the identification of highly irreplaceable areas, as prioritization softwares, such as Marxan, aim to solve the minimum set problem, i.e., meeting all targets minimizing the cost (McDonnell et al., 2002), which in the context of KBA could lead to a preference selection of PUs on coasts. This is amplified by the small representation targets, which strongly incentivize the software to select low-cost marginal PUs that contribute to reach the target without overshooting it. To overcome this issue, we suggest running prioritization analyses for KBA Criterion E using unitary cost when using softwares that aim to solve the minimum set problem. This would allow a selection of PUs without the constraint of land extent, while maintaining very highly irreplaceable PUs, as their selection is not influenced by the cost. Interestingly, this problem seems not to affect the replacement cost, although, as per KBA guidelines, we assigned the cost as land extent and set the algorithm to solve the minimum set problem.

### Recommendations for Species Distribution Modeling for KBA Criterion E applications

SDMs are a promising tool for application of KBA Criterion E, allowing assessments of many species that could be modeled at different scales (from local to continental). That said, KBA Criterion E, under the current settings, is mostly guided by the presence of range-restricted species, which strongly drive the selection of highly irreplaceable areas. This led us to the following considerations when using SDMs for KBA assessments under Criterion E. First, if the representation targets are set to the entire distribution of the species, every PUs where at least one suitable cell occurs have to be selected in order to reach the target. Thus, if the predicted suitable habitat is concentrated in a few locations, some contiguous PUs will be selected, but if the model overpredicts, even on a few cells far from the expected suitable habitat, the PUs where those cells occur will be selected as highly irreplaceable areas too. Such an issue could stem from the binarization of continuous probability outputs, which could fail to identify realistic suitable habitat, especially under different environmental conditions (Santini et al., 2021), and could introduce uncertainty in prioritization analysis (Tulloch et al., 2016). Instead of using binary habitat maps, other methods could be employed. For example, semi-binary approaches convert all values below the threshold used to binarize to zero, while retaining all others at their original values (Domisch et al., 2019), or just using the SDMs probability outputs.

Second, in the absence of clearly established spatial constraints (e.g. delimitated geographic distribution ranges) SDMs might generate large distributions of potential suitable habitat, reducing the identification of highly irreplaceable areas. In case of large distributions, the representation targets will likely be small (i.e., 10% of the suitable habitat), therefore just a few PUs will be enough to meet them, especially if many species overlap and just few PUs contribute to the representation of many features. Additionally, as many options are available to reach the targets, the algorithm will prefer selecting areas with low cost if it is set to solve the minimum set problem. To overcome this issue, SDMs used for application of KBA Criterion E should pay attention to methodological choices to handle overpredictions, such as constraining SDM predictions based on actual occurrences or expert range maps, improving variable selection (Mancini et al., 2025) or including biological interactions (Giannini et al., 2013). SDMs applications in conservation biology are continuingly increasing (Zurell et al., 2023) and they will likely be more often used also for spatial prioritization analysis. Thus we suggest KBA guidelines to include SDMs as potential source of data to represent biodiversity features and also to proactively provide some suggestions for development of SDMs explicitly focusing on KBA Criterion E applications, as already done on IUCN Red List guidelines for climate change extinction risk assessments (IUCN Standards and Petitions Committee, 2024).

Key Biodiversity Areas were developed through an extensive global consultation process. The criteria and quantitative thresholds were designed to ensure objectivity, transparency, and repeatability, drawing on technical workshops and expert inputs. Nonetheless, the methodology is not without limitations, and further testing across taxa and ecosystems may warrant future revisions. In particular, the synergy between the KBA framework and Spatial Conservation Prioritization can enhance the identification of important areas for biodiversity, helping reach the goals set in international strategic plans (i.e., Kunming-Montreal Global Biodiversity Framework), and tackling the current biodiversity crisis. KBA Criterion E could play a crucial role in integrating these two approaches, therefore its accuracy and consistency should be improved.

## Acknowledgements

G.M. and M.D.M acknowledge support from the project GaP, funded by Biodiversa+, the European Biodiversity Partnership under the 2021–2022 BiodivProtect joint call for research proposals, co-funded by the European Commission (GA N°101052342) and the Italian Ministry of University and Research (CUP B83C23001960001). M.C., W.T., and M.T. received funding from the European Union’s Horizon Europe under grant agreement N° 101060429 (NaturaConnect).

## Notes

### Competing Interest Statement

The authors have declared no competing interest.

## References

Araújo, M. B., Alagador, D., Cabeza, M., Nogués-Bravo, D., & Thuiller, W. (2011). Climate change threatens European conservation areas. Ecology Letters, 14(5), 484–492. 10.1111/j.1461-0248.2011.01610.x

Ardron, J. A., Possingham, H. P., & Klein, C. J. (2010). Marxan Good Practices Handbook (Version 2). Pacific Marine Analysis and Research Association. https://www.pacmara.org

Ball, I. R., Possingham, H., & Watts, M. E. (2009). Marxan and relatives: Software for spatial conservation prioritization. Oxford University Press.

Barrett, T., Dowle, M., Srinivasan, A., Gorecki, J., Chirico, M., & Hocking, T. (2024). data.table: Extension of ‘data.frame’. https://CRAN.R-project.org/package=data.table

BirdLife International. (2025a). Important Bird Areas. https://datazone.birdlife.org/topics/important-sites-for-birds

BirdLife International. (2025b). The World Database of Key Biodiversity Areas. Developed by the KBA Partnership: BirdLife International, International Union for the Conservation of Nature, Amphibian Survival Alliance, Conservation International, Critical Ecosystem Partnership Fund, Global Environment Facility, Re:wild, NatureServe, Rainforest Trust, Royal Society for the Protection of Birds, Wildlife Conservation Society and World Wildlife Fund. Available at https://www.keybiodiversityareas.org. [Accessed (24/07/2025)].

Brun, P., Zimmermann, N. E., Hari, C., Pellissier, L., & Karger, D. N. (2022). Global climate-related predictors at kilometer resolution for the past and future. Earth System Science Data, 14(12), 5573–5603. 10.5194/essd-14-5573-2022

Cabeza, M., & Moilanen, A. (2006). Replacement cost: A practical measure of site value for cost-effective reserve planning. Biological Conservation, 132(3), 336–342. 10.1016/j.biocon.2006.04.025

Carwardine, J., Wilson, K. A., Hajkowicz, S. A., Smith, R. J., Klein, C. J., Watts, M., & Possingham, H. P. (2010). Conservation Planning when Costs Are Uncertain. Conservation Biology, 24(6), 1529–1537. 10.1111/j.1523-1739.2010.01535.x

Chamberlain, S. A., & Boettiger, C. (2017). R Python, and Ruby clients for GBIF species occurrence data (No. e3304v1). PeerJ Inc. 10.7287/peerj.preprints.3304v1

Chauvier-Mendes, Y., Pollock, L. J., Verburg, P. H., Karger, D. N., Pellissier, L., Lavergne, S., Zimmermann, N. E., & Thuiller, W. (2024). Transnational conservation to anticipate future plant shifts in Europe. Nature Ecology & Evolution, 8(3), 454–466. 10.1038/s41559-023-02287-3

Cox, N., Chanson, J., & Stuart, S. (2006). The Status and Distribution of Reptiles and Amphibians of the Mediterranean Basin. IUCN.

Di Marco, M., Brooks, T., Cuttelod, A., Fishpool, L. D. C., Rondinini, C., Smith, R. J., Bennun, L., Butchart, S. H. M., Ferrier, S., Foppen, R. P. B., Joppa, L., Juffe-Bignoli, D., Knight, A. T., Lamoreux, J. F., Langhammer, P. F., May, I., Possingham, H. P., Visconti, P., Watson, J. E. M., & Woodley, S. (2016). Quantifying the relative irreplaceability of important bird and biodiversity areas. Conservation Biology, 30(2), 392–402. 10.1111/cobi.12609

Di Marco, M., Watson, J. E. M., Possingham, H. P., & Venter, O. (2017). Limitations and trade-offs in the use of species distribution maps for protected area planning. Journal of Applied Ecology, 54(2), 402–411. 10.1111/1365-2664.12771

Domisch, S., Friedrichs, M., Hein, T., Borgwardt, F., Wetzig, A., Jähnig, S. C., & Langhans, S. D. (2019). Spatially explicit species distribution models: A missed opportunity in conservation planning? Diversity and Distributions, 25(5), 758–769. 10.1111/ddi.12891

Farooq, H., Antonelli, A., & Faurby, S. (2023). A call for improving the Key Biodiversity Areas framework. Perspectives in Ecology and Conservation, 21(1), 85–91. 10.1016/j.pecon.2023.02.002

Ferrier, S., Pressey, R. L., & Barrett, T. W. (2000). A new predictor of the irreplaceability of areas for achieving a conservation goal, its application to real-world planning, and a research agenda for further refinement. Biological Conservation, 93(3), 303–325. 10.1016/S0006-3207(99)00149-4

GBIF.org. (2023). Occurrence Download. 10.15468/dl.wzfhjd; 10.15468/dl.trbz6u; 10.15468/dl.nmbycd

Giannini, T. C., Chapman, D. S., Saraiva, A. M., Alves-dos-Santos, I., & Biesmeijer, J. C. (2013). Improving species distribution models using biotic interactions: A case study of parasites, pollinators and plants. Ecography, 36(6), 649–656. 10.1111/j.1600-0587.2012.07191.x

Guisan, A., Thuiller, W., & Zimmermann, N. E. (2017). Habitat Suitability and Distribution Models: With Applications in R. Cambridge University Press.

Guisan, A., Tingley, R., Baumgartner, J. B., Naujokaitis-Lewis, I., Sutcliffe, P. R., Tulloch, A. I. T., Regan, T. J., Brotons, L., McDonald-Madden, E., Mantyka-Pringle, C., Martin, T. G., Rhodes, J. R., Maggini, R., Setterfield, S. A., Elith, J., Schwartz, M. W., Wintle, B. A., Broennimann, O., Austin, M., … Buckley, Y. M. (2013). Predicting species distributions for conservation decisions. Ecology Letters, 16(12), 1424–1435. 10.1111/ele.12189

Gurobi Optimization, L. (2024). Gurobi Optimizer Reference Manual. https://www.gurobi.com

Hanson, J. O., Schuster, R., Strimas-Mackey, M., Morrell, N., Edwards, B. P. M., Arcese, P., Bennett, J. R., & Possingham, H. P. (2024). Systematic conservation prioritization with the prioritizr R package. Conservation Biology, e14376. 10.1111/cobi.14376

Hijmans, R. J. (2024). terra: Spatial Data Analysis. https://CRAN.R-project.org/package=terra

IUCN. (2016). A Global Standard for the Identification of Key Biodiversity Areas. Version 1.0. First edition. Gland, Switzerland: IUCN.

IUCN. (2023). The IUCN Red List of Threatened Species. Version 2023-1. https://www.iucnredlist.org

IUCN Standards and Petitions Committee. (2024). Guidelines for Using the IUCN Red List Categories and Criteria. Version 16. https://nc.iucnredlist.org/redlist/content/attachment_files/RedListGuidelines.pdf

Karger, D. N., Conrad, O., Böhner, J., Kawohl, T., Kreft, H., Soria-Auza, R. W., Zimmermann, N. E., Linder, H. P., & Kessler, M. (2017). Climatologies at high resolution for the earth’s land surface areas. Scientific Data, 4(1), 170122. 10.1038/sdata.2017.122

KBA Standards and Appeals Committee of IUCN SSC/WCPA. (2022). Guidelines for using A Global Standard for the Identification of Key Biodiversity Areas. Version 1.2. Gland, Switzerland: IUCN. 10.2305/IUCN.CH.2022.KBA.1.2.en

Kenyeres, Z., Rácz, I. A., & Varga, Z. (2009). Endemism hot spots, core areas and disjunctions in European Orthoptera. Acta Zoologica Cracoviensia - Series B: Invertebrata, 52(1–2), 189–211. 10.3409/azc.52b_1-2.189-211

Körner, C. (2004). Mountain Biodiversity, Its Causes and Function. AMBIO: A Journal of the Human Environment, 33(sp13), 11–17. 10.1007/0044-7447-33.sp13.11

Kukkala, A. S., & Moilanen, A. (2013). Core concepts of spatial prioritisation in systematic conservation planning. Biological Reviews, 88(2), 443–464. 10.1111/brv.12008

Kullberg, P., Di Minin, E., & Moilanen, A. (2019). Using key biodiversity areas to guide effective expansion of the global protected area network. Global Ecology and Conservation, 20, e00768. 10.1016/j.gecco.2019.e00768

Mancini, G., Di Marco, M., Carboni, M., Cerretti, P., Maiorano, L., & Santini, L. (2025). On the Importance of Expert-Informed Variable Selection in Species Distribution Modelling. Journal of Biogeography, e70037. 10.1111/jbi.70037

Mancini, G., Santini, L., Cazalis, V., Akçakaya, H. R., Lucas, P. M., Brooks, T. M., Foden, W., & Di Marco, M. (2024). A standard approach for including climate change responses in IUCN Red List assessments. Conservation Biology, e14227. 10.1111/cobi.14227

Margules, C. R., & Pressey, R. L. (2000). Systematic conservation planning. Nature, 405(6783), 243–253. 10.1038/35012251

McDonnell, M. D., Possingham, H. P., Ball, I. R., & Cousins, E. A. (2002). Mathematical Methods for Spatially Cohesive Reserve Design. Environmental Modeling & Assessment, 7(2), 107–114. 10.1023/A:1015649716111

Moilanen, A., & Arponen, A. (2011). Setting conservation targets under budgetary constraints. Biological Conservation, 144(1), 650–653. 10.1016/j.biocon.2010.09.006

Moilanen, A., Wilson, K., & Possingham, H. (2009). Spatial Conservation Prioritization: Quantitative Methods and Computational Tools. Oxford University Press. https://www.oup.com.au/books/others/9780199547760-spatial-conservation-prioritization

Myers, N., Mittermeier, R. A., Mittermeier, C. G., da Fonseca, G. A. B., & Kent, J. (2000). Biodiversity hotspots for conservation priorities. Nature, 403(6772), 853–858. 10.1038/35002501

Pedersen, T. L. (2023). patchwork: The Composer of Plots. R package version 1.1.3. https://CRAN.R-project.org/package=patchwork

Poggio, L., de Sousa, L. M., Batjes, N. H., Heuvelink, G. B. M., Kempen, B., Ribeiro, E., & Rossiter, D. (2021). SoilGrids 2.0: Producing soil information for the globe with quantified spatial uncertainty. SOIL, 7(1), 217–240. 10.5194/soil-7-217-2021

Posit Team. (2023). RStudio: Integrated Development Environment for R.

Pressey, R. L., Johnson, I. R., & Wilson, P. D. (1994). Shades of irreplaceability: Towards a measure of the contribution of sites to a reservation goal. Biodiversity & Conservation, 3(3), 242–262. 10.1007/BF00055941

R Core Team. (2025). R: A language and environment for statistical computing. In R Foundation for Statistical Computing, Vienna, Austria. URL https://www.R-project.org/.

Sandström, E., Namasivayam, A., Oostdijk, S., Scherpenhuijzen, N., Debonne, N., & Verburg, P. H. (2023). Land system map for Europe (Version 6) [Dataset]. DataverseNL. 10.34894/THARMK

Santini, L., Benítez-López, A., Maiorano, L., Čengić, M., & Huijbregts, M. A. J. (2021). Assessing the reliability of species distribution projections in climate change research. Diversity and Distributions, 27(6), 1035–1050. 10.1111/ddi.13252

Serra, N., Kockel, A., Game, E. T., Grantham, H., Possingham, H. P., & McGowan, J. (2020). Marxan User Manual: For Marxan version 2.43 and above. The Nature Conservancy (TNC), Arlington, Virginia, United States and Pacific Marine Analysis and Research Association (PacMARA).

Si-Moussi, S., & Thuiller, W. (2024). Species habitat suitability of European terrestrial vertebrates for contemporary climate and land use (Version 1) [Dataset]. German Centre for Integrative Biodiversity Research. 10.25829/wpfn43

Smith, R. J., Bennun, L., Brooks, T. M., Butchart, S. H. M., Cuttelod, A., Di Marco, M., Ferrier, S., Fishpool, L. D. C., Joppa, L., Juffe-Bignoli, D., Knight, A. T., Lamoreux, J. F., Langhammer, P., Possingham, H. P., Rondinini, C., Visconti, P., Watson, J. E. M., Woodley, S., Boitani, L., … Scaramuzza, C. A. de M. (2019). Synergies between the key biodiversity area and systematic conservation planning approaches. Conservation Letters, 12(1), e12625. 10.1111/conl.12625

Thuiller, W. (2024). Ecological niche modelling. Current Biology, 34(6), R225–R229. 10.1016/j.cub.2024.02.018

Thuiller, W., Lavergne, S., Roquet, C., Boulangeat, I., Lafourcade, B., & Araujo, M. B. (2011). Consequences of climate change on the tree of life in Europe. Nature, 470(7335), 531–534. 10.1038/nature09705

Troudet, J., Grandcolas, P., Blin, A., Vignes-Lebbe, R., & Legendre, F. (2017). Taxonomic bias in biodiversity data and societal preferences. Scientific Reports, 7(1), 1–14. 10.1038/s41598-017-09084-6

Tulloch, A. I. T., Sutcliffe, P., Naujokaitis-Lewis, I., Tingley, R., Brotons, L., Ferraz, K. M. P. M. B., Possingham, H., Guisan, A., & Rhodes, J. R. (2016). Conservation planners tend to ignore improved accuracy of modelled species distributions to focus on multiple threats and ecological processes. Biological Conservation, 199, 157–171. 10.1016/j.biocon.2016.04.023

Tzivanopoulos, M., Gallien, L., Münkemüller, T., Hedde, M., Michez, D., de Manincor, N., Fontaine, C., Gérard, S., Sentil, A., & Thuiller, W. (in review). Climate dominance modulated by soil and land-use: Multi-guild drivers of European invertebrate distributions.

Watson, J. E. M., Venegas-Li, R., Grantham, H., Dudley, N., Stolton, S., Rao, M., Woodley, S., Hockings, M., Burkart, K., Simmonds, J. S., Sonter, L. J., Sreekar, R., Possingham, H. P., & Ward, M. (2023). Priorities for protected area expansion so nations can meet their Kunming-Montreal Global Biodiversity Framework commitments. Integrative Conservation, 2(3), 140–155. 10.1002/inc3.24

Wickham, H., Averick, M., Bryan, J., Chang, W., McGowan, L., François, R., Grolemund, G., Hayes, A., Henry, L., Hester, J., Kuhn, M., Pedersen, T., Miller, E., Bache, S., Müller, K., Ooms, J., Robinson, D., Seidel, D., Spinu, V., … Yutani, H. (2019). Welcome to the Tidyverse. Journal of Open Source Software, 4(43), 1686. 10.21105/joss.01686

Zizka, A., Silvestro, D., Andermann, T., Azevedo, J., Duarte Ritter, C., Edler, D., Farooq, H., Herdean, A., Ariza, M., Scharn, R., Svantesson, S., Wengström, N., Zizka, V., & Antonelli, A. (2019). CoordinateCleaner: Standardized cleaning of occurrence records from biological collection databases. Methods in Ecology and Evolution, 10(5), 744–751. 10.1111/2041-210X.13152

Zurell, D., Fritz, S. A., Rönnfeldt, A., & Steinbauer, M. J. (2023). Predicting extinctions with species distribution models. Cambridge Prisms: Extinction, 1, e8. 10.1017/ext.2023.5

