## Appendix S2 for "Opportunities and challenges for applying Key Biodiversity Areas Criterion E at large spatial scales"

Supplementary Materials

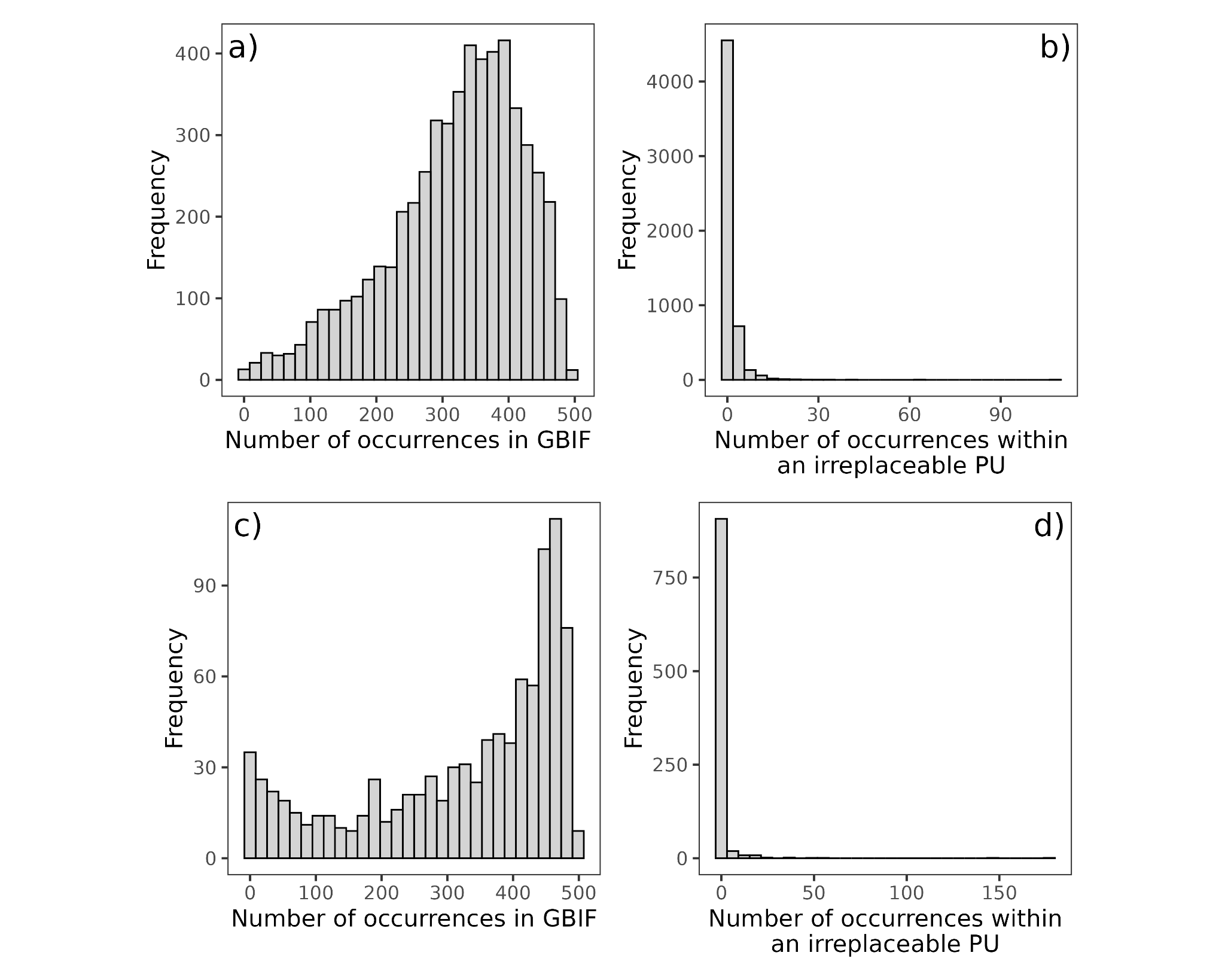

Fig. S1. Histograms of overall GBIF occurrences for insects (a) and tetrapods (c), with histograms representing GBIF occurrences within irreplaceable planning units selected using the selection frequency at 10 km resolution for insects (b) and tetrapods (d).

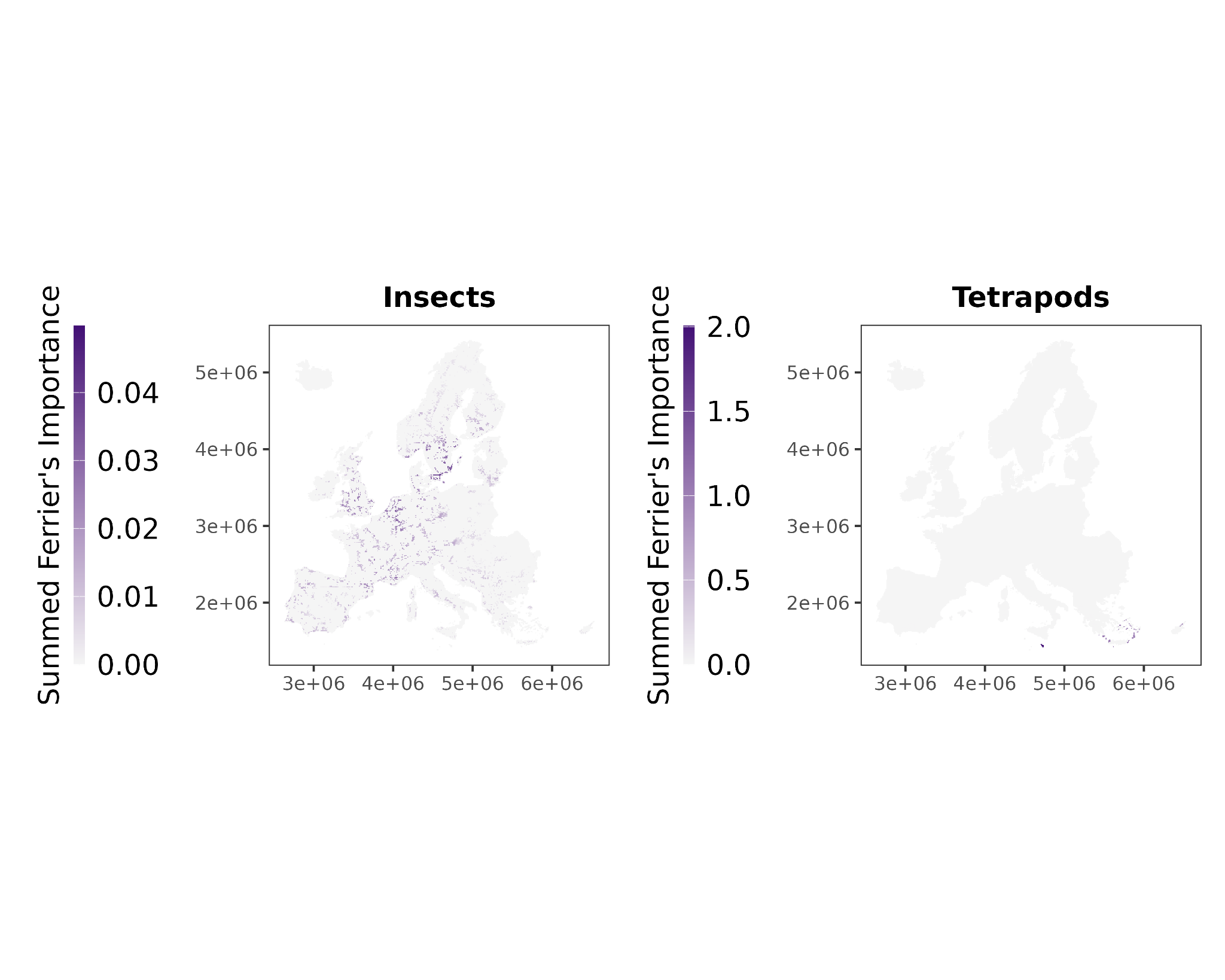

Fig. S2. Geographical pattern of Summed Ferrier’s importance values for insects and tetrapods at 10 km resolution.

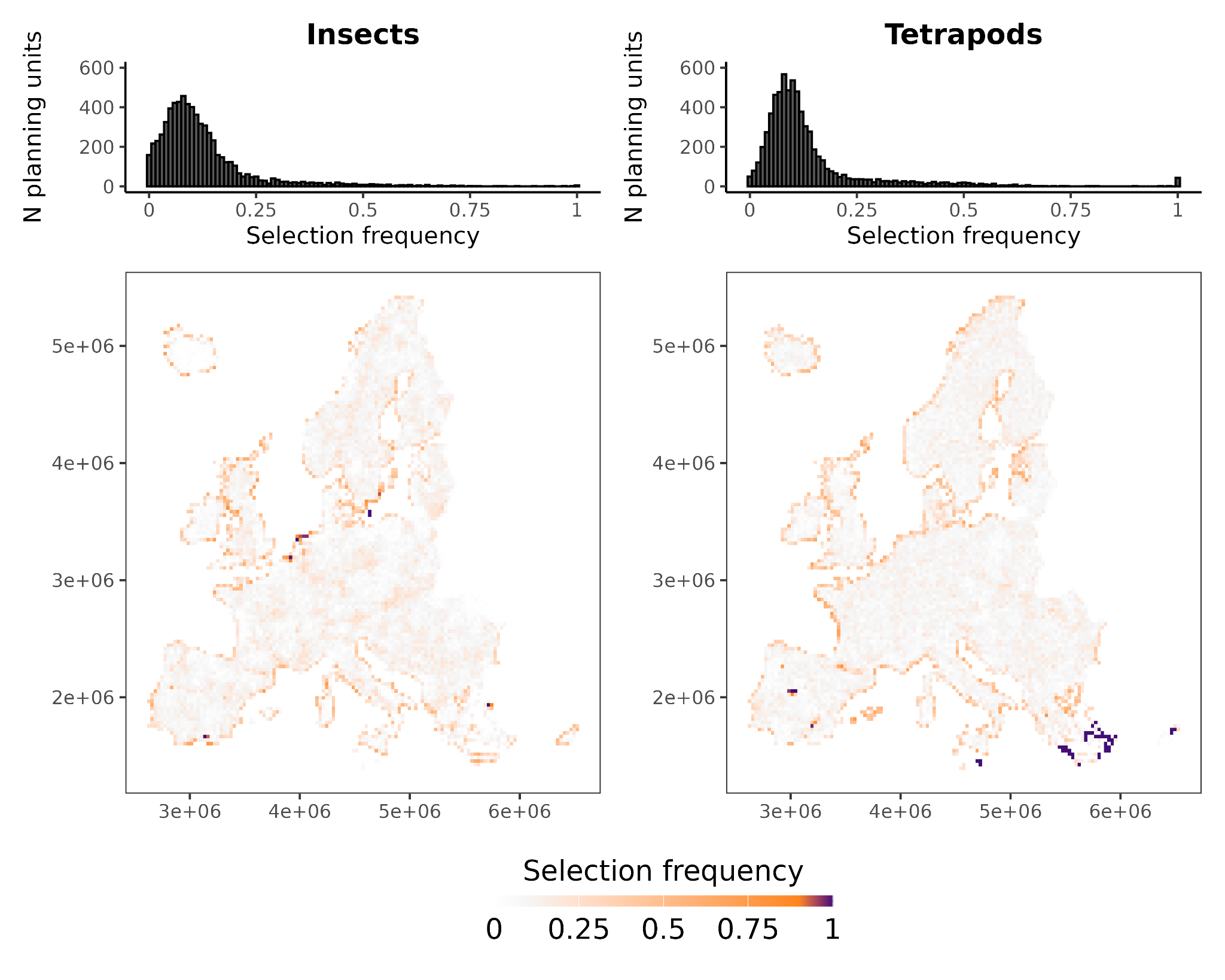

Fig. S3. Selection frequency (SF) values for insects and tetrapods at 30 km resolution. Maps represent the geographical pattern of SF values for insects and tetrapods. Histograms represent the distribution of SF values for insects and tetrapods.

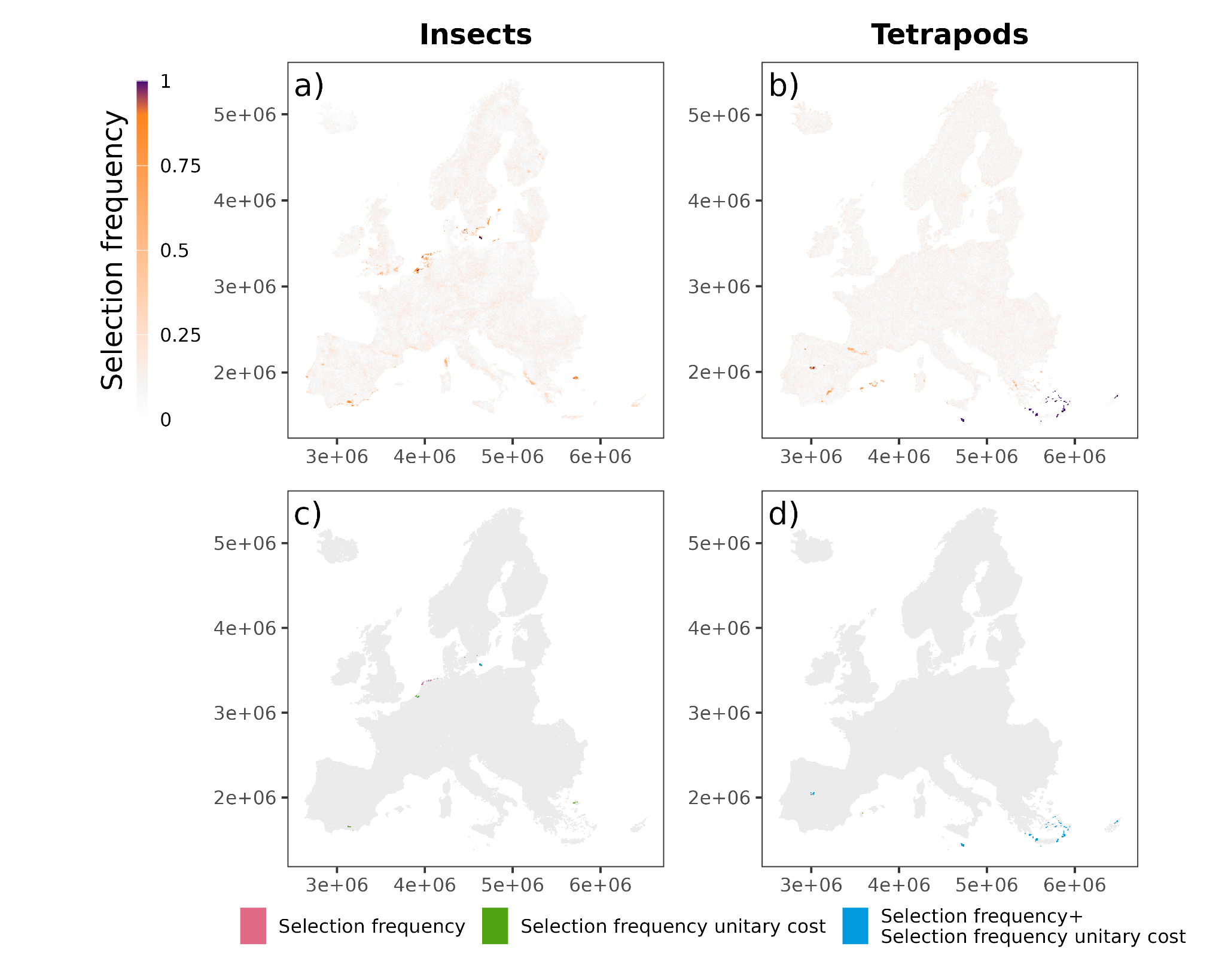

Fig. S4. Selection frequency at 10 km resolution using unitary cost for insects (a) and tetrapods (b). Highly irreplaceable planning units identified using standard settings for cost (where each PU has a cost representing the extent of land in that PU; red), with dummy unitary cost (green) or common to the two analyses (blue) for insects (c) and tetrapods (d).

Tab. S1. Estimates of European insect species in Europe.

| **Order** | **Group** | **N species analyzed** | **N species in Europe** | **Reference** |
| --- | --- | --- | --- | --- |
| Hymenoptera | Anthophila | 475 | 2138 | Ghisbain et al., 2023, Zootaxa |
|  | Ichneumonoidea | 200 | 5865 | van Achterberg et al., 2017, Biodiversity Data Journal |
|  | Formicidae | 108 | 622 | Borowiec 2014, Genus - Monograph |
| Coleoptera | Carabidae | 442 | 2700 | Capinera, J. L. (Ed.) 2008, Encyclopedia of Entomology |
| Lepidoptera | Heterocera | 3159 | 10000 | European Red List of butterflies |
|  | Rhopalocera | 344 | 576 | Van Swaay, C.A.M. & Warren, M.S. 1999, Red Data book of European butterflies (Rhopalocera) |
| Odonata |  | 115 | 146 | European Red List of Dragonflies & Damselflies (Odonata) |
| Orthoptera |  | 276 | 1229 | Ivković et al. 2024, ARTICULATA Beiheft |
| Diptera | Syrphidae | 410 | 951 | Speight 2020, Species accounts of European Syrphidae |

**References**

Borowiec, L. (2014). Catalogue of ants of Europe, the Mediterranean Basin and adjacent regions (Hymenoptera: Formicidae). Genus (Wrocław)*, 25*(1-2), 1-340.

Capinera, J. L. (Ed.). (2008). *Encyclopedia of entomology*. Springer Science & Business Media.

European Environment Agency. (2025). *Protecting and restoring Europe's wild pollinators and their habitats*. European Environment Agency. https://www.eea.europa.eu/en/newsroom/news/protecting-and-restoring-europes-wild-pollinators-and-their-habitats

Ghisbain, G., Rosa, P., Bogusch, P., Flaminio, S., Le Divelec, R., Dorchin, A., ... & Reverte, S. (2023). The new annotated checklist of the wild bees of Europe (Hymenoptera: Anthophila). *Zootaxa*, *5327*(1), 1-147.

De Knijf, G., Billqvist, M., van Grunsven, R.H.A., Prunier, F., Vinko, D., Trottet, A., Bellotto, V., Clay, J. and Allen, D.J. (2024). Measuring the pulse of European biodiversity. European

Red List of Dragonflies & Damselflies (Odonata). Brussels, Belgium: European

Commission. 46 pp.

Ivković, S., Husemann, M., Tumbrinck, J. and Heller K.J. (2024). An updated checklist of European Orthoptera. *ARTICULATA Beiheft*, 39, 1-58

Speight, M. C. (2020). Species accounts of European Syrphidae, 2020. *Syrph the net, the database of European Syrphidae (Diptera)*, *104*, 314.

van Achterberg, K., Taeger, A., Blank, S. M., Zwakhals, K., Viitasaari, M., Yu, D. S. K., & de Jong, Y. (2017). Fauna Europaea: Hymenoptera–Symphyta & Ichneumonoidea. *Biodiversity Data Journal*, (5), e14650.

van Swaay, C., Cuttelod, A., Collins, S., Maes, D., Munguira, M. L., Šašić, M., ... & Wynhoff, I. (2010). *European red list of butterflies*. Luxembourg: Publications Office of the European Union.

Van Swaay, C., & Warren, M. (1999). *Red data book of European butterflies (Rhopalocera)* (Vol. 99). Council of Europe.
